# Visualization and review of reads alignment on the graphical pan-genome with VAG

**DOI:** 10.1101/2023.01.20.524849

**Authors:** Fangping Li, Haifei Hu, Zitong Xiao, Jingming Wang, Jieying Liu, Deshu Zhao, Yu Fu, Yijun Wang, Xue Yuan, Suhong Bu, Xiaofan Zhou, Junliang Zhao, Shaokui Wang

**Author notes:** Corresponding author: Shaokui Wang, Junliang Zhao, E-mail address: Shaokui Wang, Junliang Zhao. These authors contributed equally to this work (Fangping Li, Haifei Hu, Zitong Xiao).

## Abstract

Recent advances in pangenomics have led to the rapid development of graph-based pangenomes that code genetic variant as nodes and edges preserve the contiguity of the sequence and structural variation between individuals. Data visualization is an essential component of genomic data analysis. However, the further application of the graph-based pangenome is still suffered from lacking bioinformatics tools to visualize graph format pangenomes and understand the reads alignment on graph pangenomes. In this research, we developed a novel bioinformatics platform, VAG (Visualizing read alignments in graph genomes), to overcome these challenges. VAG includes multifunctional modules integrated into a single command line and an online visualization platform supported through a web server. This tool can extract specific sequence regions from a graph pangenome and display read alignments on different paths of a graph pangenome. In addition, VAG provides population-level presence/absence variations frequency analysis and sequence path navigation to identify the population differentiation regions. To demonstrate the usage, we investigated genetic variations using a rice graph pangenome with population-level sequencing data to identify important genes and gene clusters underlying the *indica–japonica* differentiation with VAG. After investigating read alignments on the graph pangenome, we identified many false-positive alignments due to TE insertions. To reduce the impact of these misleading alignments, we developed a navigation module to determine and filter those false-positive alignments based on the pair-end alignment information. The utilization of mate-pair information in VAG provides a reliable reference for variation identification. Additionally, we developed a VAG web server to provide a user-friendly and interactive platform to visualize the read alignment data. VAG was also applied to SV discovery in the cucumber and soybean graph-based pangenome and details of VAG can be accessed by the following website (https://ricegenomichjx.xiaomy.net/VAG/sequenceextraction.php).

## Introduction

Advances in sequencing technologies and bioinformatics have driven genomics research in assembling reference genomes of plant species. The comparison among genome assemblies led researchers to realize that a genome assembled from a single individual provides an incomplete representation of the genetic landscape for the species, owing to the genetic variation observed within populations (Golicz et al., 2016; Montenegro et al., 2017). Therefore, a pangenome containing a collection of genomic sequences from a species or clade is necessary (Bayer et al., 2020). The pangenome comprises a core genome shared by all individuals, and a dispensable genome, with regions lost in one or more individuals. Recently, the graph-based pangenome emerged as an alternative to linear pangenome representations (*de novo* and iterative pangenomes). The graph-based pangenome integrates both pangenome sequences and variant information, displaying structural variations (Edwards and Batley, 2022).

Although graph pangenomes provide more potential to facilitate identifying and locating causal variants of significant agronomic traits, the broader application of the graph-based pangenome is still limited by lacking available tools to visualize structural variations and understand the read alignment on graph format pan-genomes (Wang et al., 2022). Visualization of read alignments is a key to genomic research since read alignment details are essential for detecting genetic variations and manually identifying and correcting potential alignment errors (Thorvaldsdóttir et al., 2013). Tools such as IGV, Jbrowse and Appollo were developed to visualize read alignments and genetic variations on linear format reference genomes (Skinner et al., 2009; Thorvaldsdóttir et al., 2013). Still, these tools cannot be directly applied in more complex graph format pangenomes. Recent studies have also attempted to develop different tools for visualizing graph-based pangenomes, including Bandage, ODGI, GFAviz, and the viz module of Vg (Guarracino et al., 2022; Hickey et al., 2020). However, all these tools mainly focus on assembly graph visualization, helping display genetic variations between different genome assemblies and understand the extended structure of the pangenome.

Currently, graph-based pangenome approaches are relatively new and have not been widely unlisted by the research community, which require corresponding algorithms and bioinformatics tools to fill in the gaps, especially data visualization of read alignments in a graph-based pangenome. Therefore, in this study, we introduced a novel bioinformatic platform VAG, to enhance the scientific community’s understanding of the complex graph-based pangenomes by allowing faster and instinctive visualization of read alignments and flexible interaction on graph-based pangenome datasets.

## Method and material

### The main function and dependencies of VAG

#### Total working flow

VAG was a comprehensive analysis platform for the graphical pangenome data and its corresponding reads alignment. The graphsamtools module in VAG provides a paradigm to extract the alignment information according to the specified interval. The extracted information can be visualized by command lines or interactive web-sever. VAG also integrated the modules of population-level PAV analysis with the graphical pangenome based on read alignments (Figure 1A).

**Figure 1.**
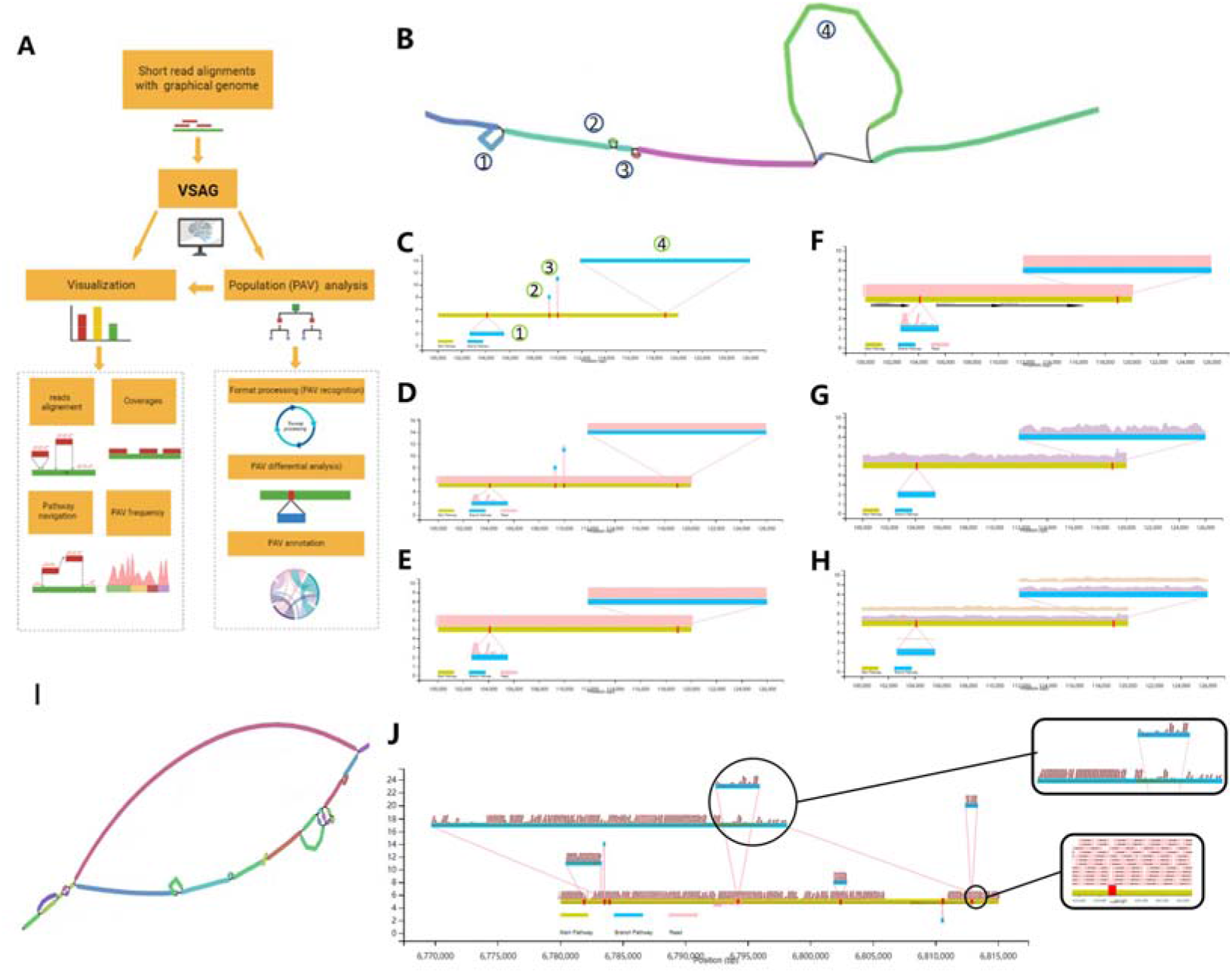
VAG allows the visualization of read alignments, path coverage of read alignments, comparison of read alignments of multiple lines and inversions in the graph-based pan-genome. **A:** Workflow and infrastructure of VAG. **B-H:** Different visualization examples of comparison of the rice graph-based pangenome. **B:** Graphical visualization in the rice graph-based pangenome using Bandage. **C:** Visualization of the same region in Figure B using VAG. **D:** Visualization of short reads alignment in this region. **E:** Branch filtering according to the sequence length and read alignment condition. **F:** Display of additional feature across the graph pangenome. **G:** Visualization of the coverages of short read sequencing alignments. **H:** Alignment from multiple samples within the same location in the graph pangenome. **I:** Visualization of interval in using Bandage. **J:** Visualization of Chr8 (6,780,000-6,815,000) using VAG

#### Paths and alignment information extraction based on reference genome interval

Based on the targeted genome interval of the main pathway or reference genome, users can easily extract and visualize the main pathway and the associated branching path from the variation graph (.gfa format). Softwares of GFAtool, Bedtool and Samtool were integrated into VAG in this process (Li et al., 2020).

#### Visualization of PAVs of the population with graph-based pangenome short fragment alignment

The re-sequencing data of 123 *japonica* and 207 *indica* rice varieties were utilized to investigate the differentiation between subspecies. The map module of Vg was used to align the short reads data from the rice population to the graphical pangenome (Hickey et al., 2020). For functional gene analysis, we selected the longest transcript of each gene as the representative transcript.

#### Path navigation based on pair-end information

In this step, the SupportSQ module restores each path in the variation graph to the segment form (Sx……Sx+i) generated by Minigraph based on the branch information (Li et al., 2020). The SupportSQ module further sorted the segment according to the start points of branches on the main path. The first segment from the reference genome is set as the start point and assumed to be the reliable segment. The other reliable path is extended and labelled based on whether the two ends contain pair-end information linked with the previous reliable path. If the breakpoint of a reliable path cannot be extended, VAG will try to restart the reliable path on the main path by determining a read coverage threshold (including depth and length). Branch arrangements supported by mate pair information between branch nodes are classified as reliable.

### Genome sequencing and de novo assembly and genome data download

Young leaves from two widely cultivated *indica* varieties, IND-91 and IND-86, were used for DNA extraction. High-quality genomic DNA was extracted using the QIAGEN genome kit. DNA integrity was detected by 0.75% agarose gel electrophoresis. DNA purity and quantity were detected by Nanodrop (OD260/280 = 1.8-2.0, OD260/230 = 2.0-2.2) and Qubit, respectively. Long read sequencing data used for genome assembly were generated from PacBio sequel II platform, while the short reads data utilized in alignment and genome polished were generated from Illumina NovaSeq 6000 platform. The two rice genomes were *de novo* assembled by Software MECAT2 and polished using NextPolish (Xiao et al., 2017).

### Construction of graphical pan-genome and short-reads alignment

The rice graphical pan-genome was built using minigraph in this study. The most widely used rice reference genome, MSU7 of Nipponbare (*japonica*), was used as the backbone reference genome for generating the main pathway. Two *japonica* varieties (ZH11, DHX2), one circum-basmati group (Basmati1), and six *indica* varieties (G630, TM, WSSM, TF, HZ, and R498) were screened by phylogenetic relationships (Qin et al., 2021). Genome assembly release in previous research was utilized to construct the graph pangenome in *Cucumis sativus* (cucumber) and *Glycine max* (Soybean) (Liu et al., 2020; Li et al., 2022). The alignment of sequencing read and variation detection to the graph pangenome was performed by vg map and vg giraffe (Hickey et al., 2020; Sirén, et al., 2021). Bandage and VAG were utilized to visualize the graph and read alignments, respectively (Wick et al., 2015).

### Data available

The example databases and platforms can be visited and browsed through the website https://ricegenomichjx.xiaomy.net/VAG/sequenceextraction.php and the VAG is freely available at Github (https://github.com/lipingfangs/VAG).

## Result

### Construction of a graphic pan-genome and short reads alignments

For a comprehensive test and display of VAG, a rice graph-based pan-genome was constructed to indicate the usage of VAG and demonstrate how VAG can help understand the short read alignments in a graph-based pangenome and dissect genetic variations on rice genomes. This rice graph-based pangenome was built using ten published high-quality rice genome assemblies with a genome size of 567.98 Mb. The constructed pangenome contained more than 160,000 segments, including a 372 Mb backbone reference genome (MSU) and 190 Mb branch path sequences absent in the reference genome. We further used GFAtool to convert the graph-based pangenome into a readable path format and used the graphann module of the VAG to merge and coordinate the annotation information from the ten rice genomes. In total, we detected 44,832 genes in our rice graph pangenome, including 7,321 novel genes present in at least one of nine rice genomes used for the graph pangenome construction but absent in the MSU backbone reference genome.

### Joint visualization of genome paths and alignments

Visualization of read alignments is an essential first step to identify genetic variations and detect potential errors in read alignments. Compared with read alignment visualization tools (e.g. IGV and Jbrowse) used for the linear format genome, handly bioinformatics tools specifically designed for visualization of read alignments on graph-based pangenome are still lacking. To overcome this challenge, we developed VAG and provided a set of handy and easily manipulative functions to visualize pangenome graphs and short read alignments. Using the VAG, we can locate and extract subregions from the rice graph pangenome using reference genome-based coordinates, achieving a similar overview of the graph-based pangenome as Bandage. For example, a 20 kb interval of Chr10 in MSU with ~17 kb of insertion branch was utilized to display (Figure 1B-C). Besides, VAG can provide details of read alignments in the graph-based pangenome that are missing in other graph-based pangenome visualization tools such as ODGI, sequenceTubeMap and Bandage (Figure 1B). Using the ShowRead module, VAG enables visualization of the short read sequencing alignment and provides read coverage information across the extracted subregion (Figure 1D-G). For handing the rice read alignments data with ~50x depth using default parameters, VAG takes about 2 minutes to visualize short read sequence alignment on the rice graph pangenome within a 50 kb sequence region in the reference path and 20 kb sequence regions in branch paths. Moreover, VAG can also display the details of gene location and additional genetic structure features (such as gene annotation and TE annotation) on the graph-based pangenome (Figure 1F). Using the Mutsamples module, VAG can visually display the coverage of read alignments of multiple lines at the same scale on the graph-based pangenome (Figure 1H). In addition, VAG enables researchers to display inversions of the graph pangenome and the direction of read alignments on the forward or reverse strands. For example, we extracted a 35 kb interval in Chr8 in the rice graph pangenome that contained an inversion. The VAG can clearly visualize this inversion indicated by two reverse arrows, while Bandage can only display it as a branch path without identifying it as an inversion (Figure 1 I-J).

### Population PAV frequency detection, variation determination, annotation, and visualization in the graphical pangenome

Presence/absence variations (PAVs) are common in plants and one of the major factors causing genetic differences between individuals and populations. PAVs affect gene function and shape phenotypes associated with population differentiation. The development of graph-based genome alignment tools (e.g. vg map and vg giraffe) enables remapping the short read sequencing data back to the graph-based pangenome and detecting PAVs (Sirén et al., 2021). To further dissect the hidden effect of PAVs, we urgently need new tools to discover and visualize gene presence/absence for graph-based pangenome at the population scale. Here, we developed the Populationfrequency module in VAG to detect PAVs at the population level by surveying the PAV frequency of genome interval coverage and the Populationfrequenvis module for visualizing PAV frequency distribution. In order to demonstrate the implementation of the populationfrequenvis module, we mapped the short read sequencing data to the constructed rice graph-based pangenome and investigated the gene PAVs between two rice subs-populations (*indica* and *japonica*). A total of 330 accessions of rice from the core collection of Rice Diversity Panel 2 (RDP2) (McCouch et al., 2016) were utilized in this study. All the accessions were resequenced by the Illumina platform for 50x depth.

We detected and identified 160.54 Mb sequences of the rice graph-based pangenome, with a significant difference (*P* value < 1E^−5^) in the frequency of PAVs. These PAVs sequences overlapped with 7,345 and 15,103 genes in the CDS and promoters’ regions, respectively, with 6,438 genes with PAVs in both regions (Figure. 2A). The results showed that the PAVs-based PCA analysis divided the accessions into two distinct clusters.

**Figure 2.**
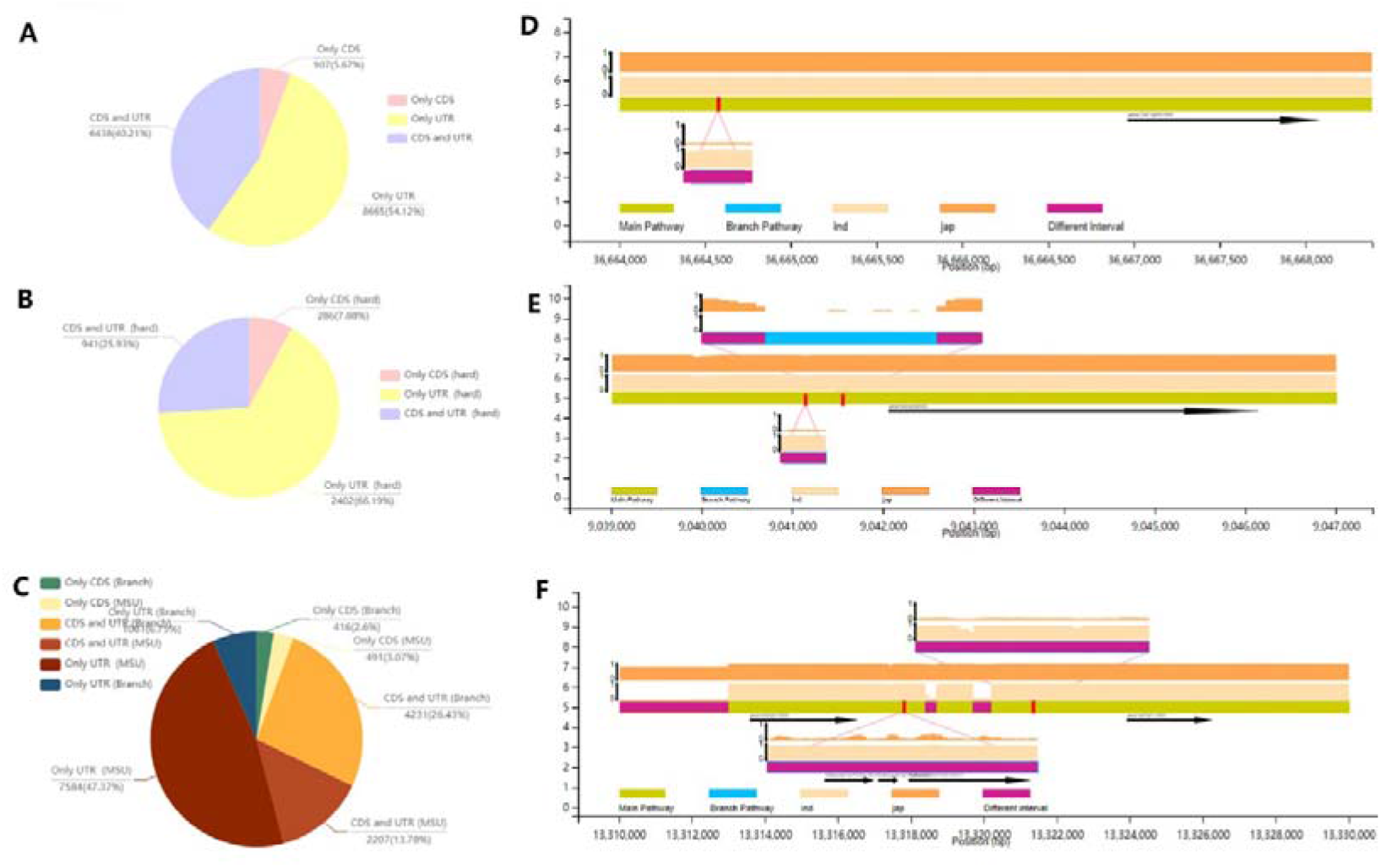
Analysis of PAV differences between *Indica* and *Japonica* rice subspecies based on short read alignments on the rice graph pangenome. **A:** The percentage of PAVs overlapped with gene functional regions (CDS and UTR- promoter). **B:** The percentage of significantly differentiated (Differential frequency > 0.8) PAVs overlapped with gene functional regions. **C:** The distribution of PAVs in different paths (Reference paths and branch paths) overlapped with gene functional regions. **D:** An example of a gene (Os01g0551600) involved in Brassinolide sterol metabolism showed significantly differentiated PAVs in the promoter region of the branch path. **E:** An example of a gene (Os02g0200200) associated with the development of spike type (TO:0000089) showed significantly differentiated PAVs in the promoter region of the branch path. **F:** An example of gene clusters showing significantly differentiated PAVs in the branch path.

From the perspective of gene structure, the PAV differentiation in the UTR regions was significantly higher than in the CDS regions (Figure 2A). It was unusual that a gene showing the PAV differentiation was only detected in CDS regions but not in UTR regions. Further investigation revealed 3,629 genes showed significantly differentiated PAVs (PAV difference frequency > 0.8) covering 26.37 Mb sequences in the rice graph-based pangenome (Figure 2B). Gene Ontology (GO) annotation of these genes showing differentiated PAVs revealed that they are functionally enriched in peptidyl-threonine modification (GO: 0018107), suggesting phosphorylation of peptidyl-threonine to form peptidyl-O-phospho-L-threonine may play a key role in *indica–japonica* differentiation. More specifically, genes with differentiated PAVs in CDS regions were enriched in 72 biological processes, including aleurone grain formation, pathogenesis, and plant hormone and energy metabolism. Genes with differentiated PAVs in UTR regions were enriched in 48 biological processes, including the metabolism of flavonoids and terpenoids, which is in line with previous studies (Deng et al., 2020). Moreover, further examination of genes with significantly differentiated PAVs (Differential frequency > 0.8) showed that they were enriched in functions associated with synthesizing steroid compounds such as brassinolide. For example, a subspecies specific PAV with a length of 963 kb was identified in the UTR of *Os01g0851600*, which was a predicted 3-oxo-5-alpha-steroid 4-dehydrogenase. This suggests that the sequence difference in steroid hormone metabolism may lead to *indica–japonica* differentiation. Besides, PAV differentiation was detected in several genes related to distinct phenotypic variations between rice subspecies. For example, a large segment of subspecies specific PAV was detected in UTR and promoter region of *OsEP3* (*Os02g0260200*), a gene reported to participate in spike type development (Borna et al., 2022), which was considered as a significant differentiation phenotype between *indica* and *japonica* (Figure 3E).

**Figure 3.**
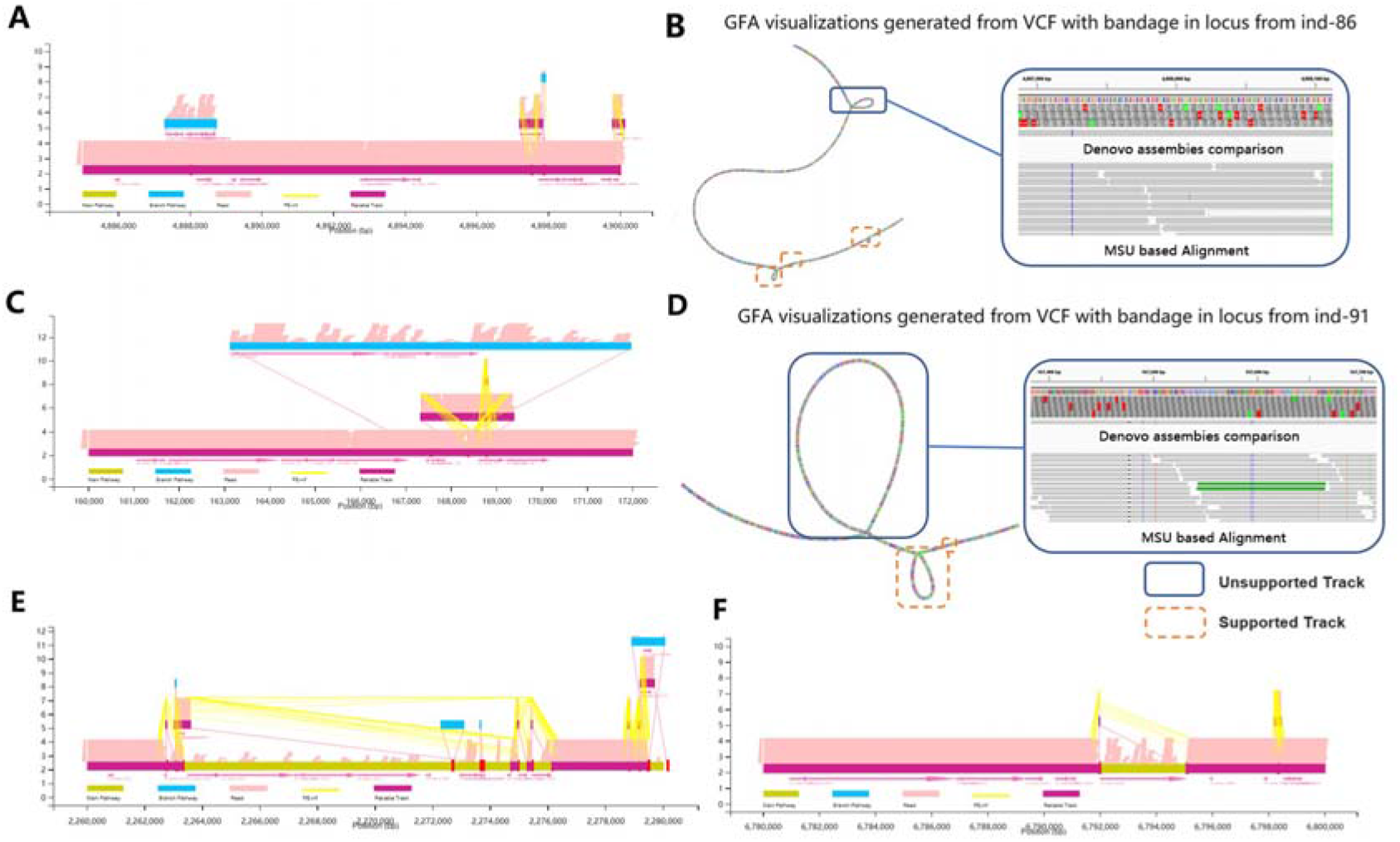
The unreliable alignments caused by transposable elements (TEs) may lead to misleading path selection in structural variations detection. **A-D:** Two examples from rice individuals IND-86 (A-B) and IND-91(C-D): The sequences in the graph-based pangenome path containing a high proportion of TEs can result in multiple incorrect short sequence alignments and lead to incorrect identification of branch paths and genetic mutations (Unreliable paths). **E:** An example showing the navigation module helps identify the unreliable paths on the reference paths.

Regarding gene distribution in the graph-based pangenome, we detected more genes with differentiated PAVs of CDS regions located in branch paths than the reference genome (MSU) paths, while a higher proportion of genes with differentiated PAVs of UTR regions were detected in the reference genome path. We also found that genes with differentiated PAVs of CDS regions were more likely located in branch paths, in which some genes completely lost whole gene sequences (Figure 2C). Meanwhile, differentiated PAVs were detected covering tandemly arrayed genes in branch paths. For example, an insertion of tandem arrayed genes was detected in an interval of 20 kb distributed in the middle of Chr7 of rice (Figure. 2F). These tandem repeat genes showed a higher presence frequency in *indica* than *japonica*, which elucidated a distinct genetic differentiation in this gene locus.

### The reviews of graph-based read alignments and potential TE interference

With the visualization of VAG, multiple alignments in sequence branches with low coverage of alignment quality were detected. To investigate this phenomenon with various sequencing data, we *de novo* assembled two representative rice genome assemblies (IND-91 and IND-86) using a combination of Pacbio long reads and Illumina short read sequencing data. From the perspective of the whole genome, the comparison indicated 50% (12,341/24,682) and 55% (13,341/25,682) variation in IND-86 and IND-91 were covered in the graph-based pan-genome. However, the alignment coverage of short sequences was detected in the remaining ~52,000 branch paths of the variation graph. This means that some branch alignments do not truly denote the sequence branch information of the individual. Both alignments examined by VAG and comparison between *de novo* assemblies indicated alignment errors occurred in this locus, leading to interference in detecting genetic variations. As a demonstration, two variations with a length of 1,017 kb and 5,035 kb were detected based on the alignment using VG for IND-86 (Chr3: 4,885,000-4,900,000) and IND-91 (Chr10: 165,000-170,000), respectively (Figure 3A-D). However, the comparison between de novo assemblies did not support these two variations. Further analysis indicated a large segment of TE distributed in the branches containing the variations. Similarly, TE interference was detected in 27,133 and 31,912 branches in the graph-based alignment of IND-91 and IND-86, respectively.

Due to the TE interference, error-prone alignments using short read sequencing can be detected in long fragment sequences in individuals that are supposed to be deletions. As a result, it is easy for software to classify one complete deletion into multiple discontinuous deletions when detecting structural variations. This error-prone SV detection may have significant implications for functional genomics research. This situation was further demonstrated in our rice graph-based pangenome. For example, compared with the MSU, the *de novo* assembled genome sequence alignment revealed a ~9 kb and ~3 kb deletion in genomes of IND-91 (Chr10: 210,000-212,000) and IND86 (Chr8: 6,780,000-6,800,000) (Figure 3E). However, short read alignments in the rice graph-based pangenome were still detected in this deletion region. This is consistent with findings in our pipeline that pair-end information does not support these error-prone alignments.

### Navigation in graphical pan-genome base on Mate Pairs information of short-reads

The use of reads with mates (mate pairs) has been applied for genome assembly to improve assemblies and generate a clear picture of the genome by resolving repeated sequences in genomic sequences. In addition, the mate pairs allowed accurate breakpoint detection and rapid structural variations (SVs) characterization. To fully utilize the graph-based pangenome, the key is to obtain a true representation of read alignments among genomic sequences, including the reference genome paths and branch paths. Graph pangenomes constructed by Minigraph preserve the coordinate of the linear reference genome, providing coordinate systems to integrate genetic segments represented in the graph pangenome.

Here, based on the single reference-based coordinate system, we developed the PairEnd module to process and display the mate pair information in VAG. The SupportSQ module in VAG was developed to navigate and understand the graph, allowing graphical identification and annotation of reliable paths supported by reads alignment with mates across different sequence segments. This module also enables users to output the consensus sequences based on reliable paths, allowing users to obtain the accurate haplotype of genome sequences. This navigation mechanism effectively avoided the error mutation mining caused by the alignment error of reads caused by TE in the multiple instance sites tested.

As a demonstration, mate pair information was not detected in the alignment in the variation related branches which were not judged as reliable by the SupportSQ module in the previous TE interference examples in IND-91(Figure 3A) and IND-86 (Figure 3C). By contrast, the branches (variations) with mate pair information determined to be reliable by supportSQ module were well supported by the comparison between *de novo* genome assemblies. This suggested VAG provides a good reference for users to judge the reliability of the path in these two intervals.

### Review the graphical based alignment to detect the accurate variation

VAG provides a paradigm for visually reviewing graph-based alignments and for more accurately displaying the population distribution and previously reported structural variations in rice. For example, *SLB1* and *SLB2* loci were reported as “deleted” in *indica* (Cardoso et al., 2014), while the phenomenon was also regarded as *japonica* regaining a fragment containing both genes, indicating an allelic relationship of this PAV was reported in this interval. However, after reviewing this fragment using VAG, we found a branch with a length of ~32 kb in *indica* and another branch with a length of ~49 kb containing SLB1 and SLB2 in *japonica* (Figure 4A), which indicates a substitution relationship rather than an allelic relationship in this locus between the *indica* and *japonica* rice subspecies. The comparison of *de novo* assemblies and mate pair information of short read alignments of IND-91 and IND-86 provide further details of sequence path of *indica* and *japonica* in the rice graph-based pangenome, which are differentiated into two types of branches (Figure 4B-E).

**Figure 4.**
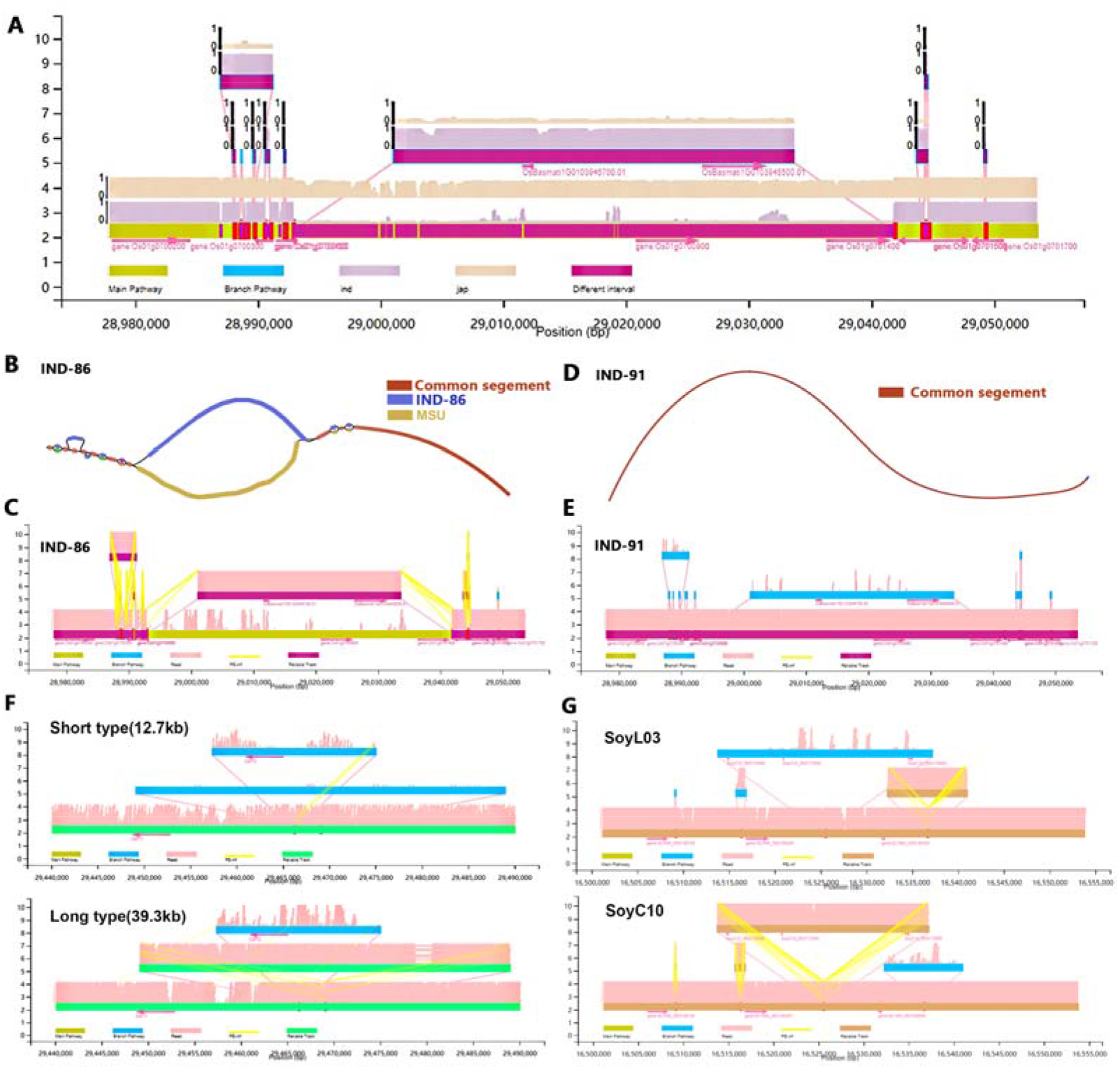
The distribution of sequences branches in the rice populationas as well as the display of individuals with the two types of branches with short reads alignment and de novo assemblies’ comparison in the SLB1/2 related interval. **A:** Significantly differentiated PAVs in branches between two types of rice sub-species. **B:** *De novo* assemblies’ comparison between MSU and IND-86 elucidated the branch’s difference. **C:** The visualization of alignment between show reads from IND-86 and graphical pan-genome, as well as mate pair information (yellow line), showed support for this variation. **D:** *De novo* assemblies’ comparison between MSU and IND-91 detected no variation. **E:** The visualization of alignment also indicated no branch difference between IND-91 and MSU (reference for pangenome). **F:** The visualization of alignment in two types of structural variations related *CaFT1* in cucumber; **G:** ~23 kb insertion from SoyC10 results in gaining of additional three genes while SoyL03 without the gene cluster contained a ~9kb insertion in the downstream of *GLYMA_06G189100*.

**Figure 5.**
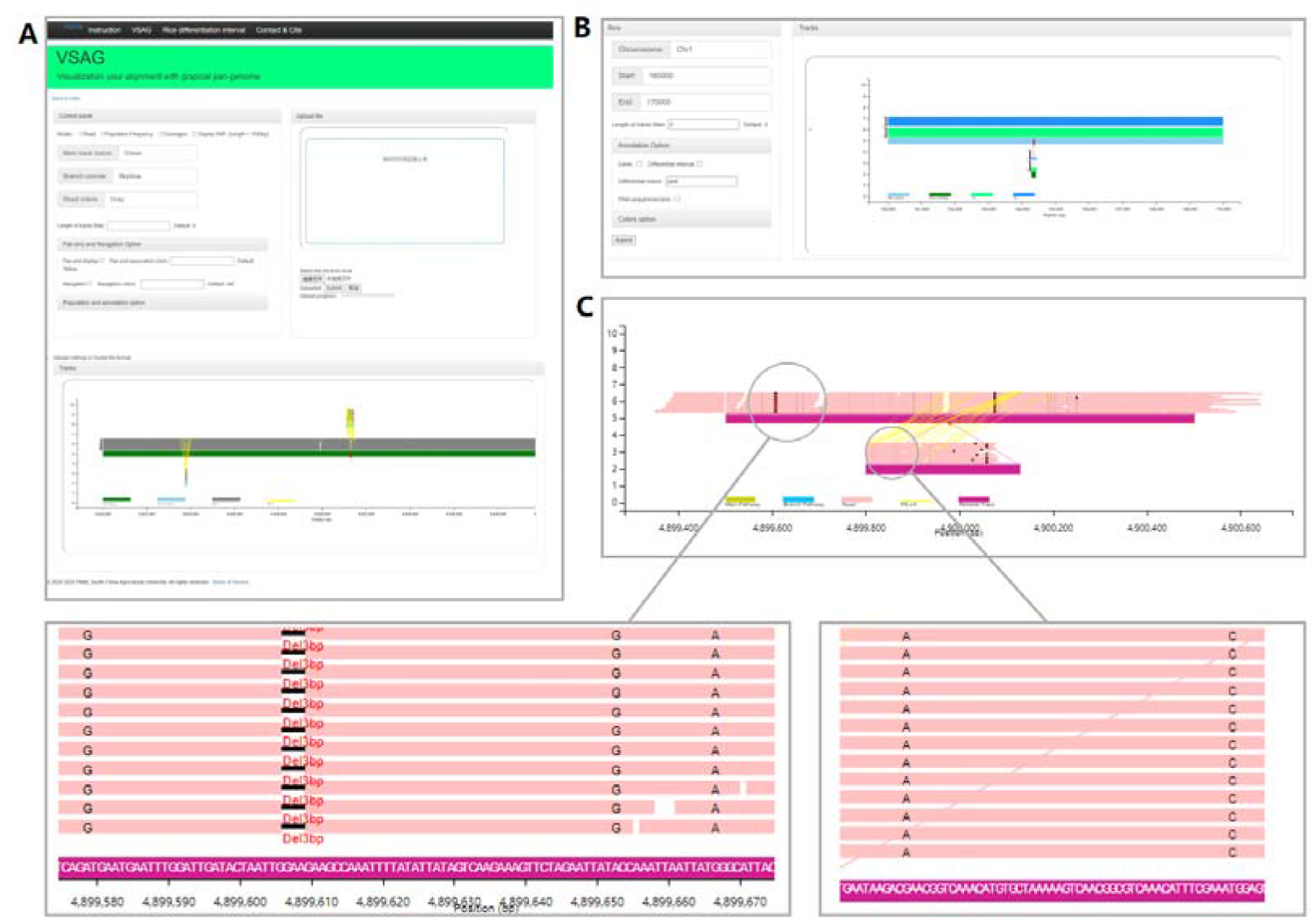
Page display and genetic variations visualization of VAG using an interactive web server. **A:** Homepage of the online VAG visualization platform. Users can use graphsamtools to extract subregions of the graph-based pangenome and perform interactive visualization through the local deployment server or our online visualization platform. **B:** Using the Webserver to visualize the PAV difference frequency between Indica and Japonica on the rice graph pangenome. Users can query the difference interval according to the physical location of the MSU reference genome. **C:** VAG can visualize SNPs and small-scale Indels (insertions/deletions) within a small segment (< 2000 bp).

In the sequence branch of *indica*, two genes were annotated, including *OsBasmati1G0103948500.01*, which is a gene highly homologous to SLB2. This suggested the subspecies specific differences caused by these two sequence branches were derived from the PAV of SLB1. Previous molecular biology and genetic studies also mainly elucidated the common loss and gain of SLB1 and SLB2, which showed obvious phenotypic differences (Cardoso et al., 2014; Qin et al., 2021). However, such gain and loss tended to be present only in SLB1. For other genes from the *Basmati* genome in this interval, the *OsBasmati1G0103946700.01* had not been reported previosouly and was detected in the sequence branch of *indica*. This gene was predicted to participate in flavonoid biosynthesis, which was in line with the enrichment result of the subspecies preference gene in rice (Figure 4C). This example shows that the review function in VAG can efficiently help us detect variations from graphical pangenome-based alignments.

Additionally, the visualization of cucumber and soybean data alignments also elucidated the universality of VAG in data from various species. As a demonstration, the visualization of alignments demonstrates the two reported structural variations were displayed in cucumber and soybean, respectively (Liu et al., 2020; Li et al., 2022). Two different complicated SVs (12.7kb (short type) / 39.3kb (long type)) related to CaFT1 were identified in the cucumber graph pangenome using VAG (Figure 4F). For soybean, the reported 23 kb insertion from SoyC10, including additional three genes, was clearly detected using VAG, while SoyL03 without this gene cluster contained an unreported insertion (~9kb) in the downstream of GLYMA_06G189100, (Figure 4G).

### Interactive display in a web-sever

To enhance the interactive ability and provide a convenient platform for users, we developed an interactive Webserver for VAG. Users can select different modules in VAG and visualize short-read sequencing alignment on the web server by uploading a dataset extracted by graphsamtools. To integrate the population level comparison and gene annotation information, users need to further upload files containing PAV frequency information or multi-population coverage information generated by graphfreq. In addition, using the SNPviz module on the web server, users can display SNPs and small INDELs (<20 bp) in small regions of the graph-based pangenome (the reference path <2,000 bp and branch paths <1,000 bp). This function provides a convenient viewer for visualizing genetic variations and helps genetic mutation mining on a panel of the graph pangenome. Besides, the server hosted by our teams also integrates the population frequency between rice subspecies with the accession utilized in this research. This provides researchers with an efficient platform to identify the subspecies differentiation interval in rice graph-based pangenome.

## Discussion

With the development of bioinformatics assembly algorisms and the falling cost of sequencing technologies, an increasing number of reference genomes are sequenced and produced, making pangenomics research that studies the genetic diversity of multiple individuals in a species become feasible. Compared with a linear format pangenome, the graph-based pangenome integrating both sequence-resolved and variant information has become a recent advance in genomic studies that facilitates identification of causal variants of phenotypic variations and genome-linked fast breeding. The graph pangenome construction tools (e.g. Minigraph and PGGB) and graph-based genome alignment tools (e.g. vg map and vg giraffe) have been applied to important crops and fruits, including bread wheat, rice, cucumber, and melon (Li et al., 2022; Vaughn et al., 2022; Bayer et al., 2022).

The current graph pangenomes visualization tools mainly focus on assembly graph visualization of different genomes. For example, Bandage (Wick et al., 2015) allows the visualization of graph-based pangenomes without further inner details to be provided and adopted. Although SequenceTubeMaps (Beyer et al., 2019) and MoMi-G (Yokoyama et al., 2019) allow the detection of structural variations from individual genomes, they are limited to providing a direct visual view of different paths of a pangenome and are unable to display population-wide read alignment information. Therefore, graph-based pangenome visualization bioinformatics tools, especially displaying the read alignments at the population level, are still lacking to help researchers further interpret the graph-based genome and dissect the biological questions hidden by the alignment data.

In this study, to fulfill the gap in data visualization of graph-based pangenomics, we developed VAG (Visualizing short-read alignment in graph genomes), with modules to analyze and visualize the read alignments on the graph-based pangenome, gene functional locus annotation and PAV frequency difference at the population level. We have identified important genes and gene clusters underlying the *indica–japonica* differentiation using a rice graph pangenome and rice short read sequencing data, which demonstrated VAG can help discover biological stories hidden behind read alignment data using a graphical pangenome.

In the meantime, read alignment in plants is complex due to large amounts of TE (Ou et al., 2018). In this study, we used VAG to analyze short read sequence alignments and compared them with *de novo* assemblies. We found that TE caused many incorrect path alignments in two rice genomes, leading to the discovery of unreliable variants. To overcome this issue, we further develop a navigation module in VAG, in which users can conduct preliminary navigation (reliable path identification) of individual local locus paths based on the read mate pairs information. Due to the complexity of the display, similar to the previously published short read alignment visualization software such as IGV (long interval choose to hide), VAG is only recommended for application for genome sequences with lengths below 100 kb. Longer sequence alignment visualizations may result in a long-running time. Lastly, we developed an interactive online platform for VAG, allowing simple and convenient visualization of genetic variations and mutation mining on the graph pangenome.

## Conclusion

VAG is a comprehensive data visualization tool for graph-based pangenomes, allowing read alignments visualization, population-wide PAV frequency difference mining and navigation of pangenome sequences based on read mate pairs. We also developed and released an easy-to-deploy Webserver, providing the interactive ability and a convenient platform for users to access the developed functions and visualize their graph pangenome data. VAG will help answer biological questions hidden behind read alignment data and facilitates the usage of graph-based pangenome for the science community.

## Acknowledgements

This study was supported by the National Natural Science Foundation of China (32072040, 31622041) and the Key Projects of Basic Research and Applied Basic Research of Guangdong Province (2019B030302006), and the Major Science and Technology Research Projects of Guangdong Laboratory for Lingnan Modern Agriculture (NT2021001), the “YouGu” Plan of Rice Research Institute of Guangdong Academy of Agricultural Sciences (2021YG001); Special Fund for Scientific Innovation Strategy-Construction of High Level Academy of Agriculture Science (R2021PY-QF002)

